# Selective predation by ants against less-honeydew secreting aphids

**DOI:** 10.1101/2024.05.02.592168

**Authors:** Takumi Matsuura, Chihiro Handa, Satobu Takahashi, Takao Itino

## Abstract

Many ant-aphid interactions illustrate the concept of a mutually beneficial association between two organisms: aphids provide ants with honeydew; in return, ants defend aphids from enemies. However, ants also often prey on the aphids they attend, and we do not know if the predation pressure causes any adaptations on the side of aphids. The aphid *Stomaphis japonica* has an obligate mutualistic association with *Lasius* ants. Here, we report evidence of selective predation of less-honeydew secreting *Stomaphis* aphids by the tending *Lasius* ants. We show that 1. the aphids are severely preyed on by the ants – up to half of the aphids’ standing population per day. 2. the ants selectively prey on the aphids which do not deliver honeydew when attacked, and 3. the frequency of aphids’ honeydew secretion is heritable. These results suggest selective predation by ants against less-honeydew secreting aphids takes place and it is heritable.

## Introduction

Many ant-aphid interactions illustrate the concept of a mutually beneficial association between two species. Ants collect honeydew excreted by aphids, often by tapping on them. The aphids, in return, benefit from protection by the ants against natural enemies, or from inadvertent hygienic consequences of ant tending [1,2].

However, ants sometimes prey on the aphids they attend [3-5]. These ant-aphid interactions are analogous to human livestock farming. The practice of ‘farming’ is also known in other organisms. For example, the nomadic ant *Dolichoderus cuspidatus* carries its mealybug livestock to suitable feeding sites [6], and Attine fungus-garden ants cultivate fungi for consumption [7]. Does such ‘livestock farming’ cause selection pressure on the target organism?

Here, we observed *Stomaphis japonica* (Aphididae) being preyed on by its attendant ant *Lasius capitatus* (Formicinae) (Fig. 1). Stable *S. japonica* colonies are always associated with *Lasius* ants, and the ants get most of their food from the aphid colonies throughout the year (Fig. 2, C.H. and T.M., personal observation). The association is thus mutually obligate for both ants and aphids. In addition, most individuals of the *S. japonica* aphids do not have wings in summer [8] (C.H. and T.M., personal observation), they held in complete captivity on ant-attending oak trees and are preyed upon by the ants. Considering the highly specific and dependent nature of this interaction, severe predation by *L. capitatus* could represent a significant source of selective pressure on *S. japonica*. So, we conducted a systematic survey of the ant-aphid interaction on *Q. acutissima* trees to know, 1. How frequently do the ants prey on the aphids? 2. Do the ants selectively prey on the less-honeydew secreting aphids? and 3. Does the frequency of honeydew secretion of aphids have a genetic basis?

**Fig. 1.**
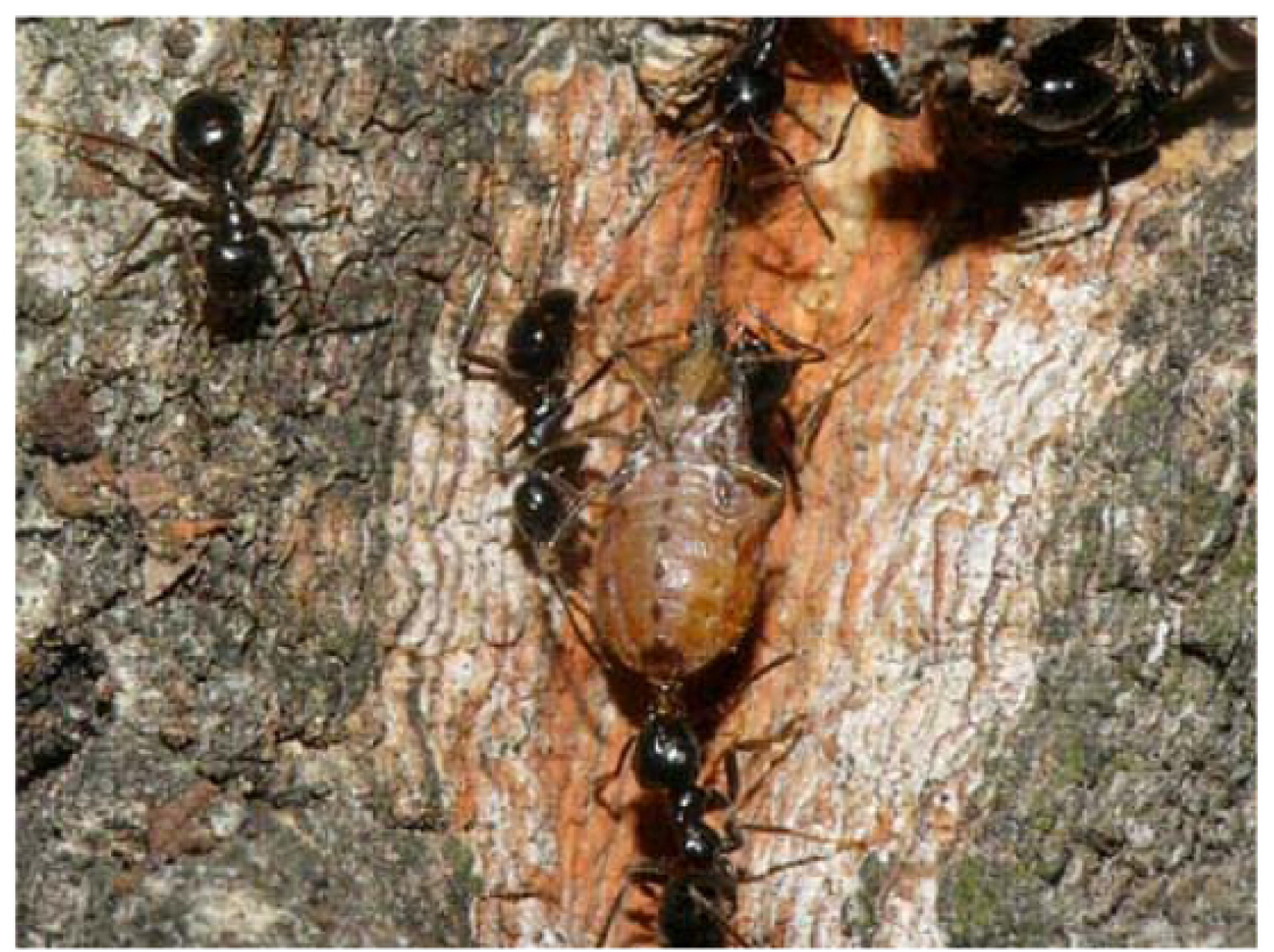
A *Stomaphis japonica* adult aphid preyed upon and transported by *Lasius capitatus* worker ants on a *Quercus acutissima* tree trunk

**Fig. 2.**
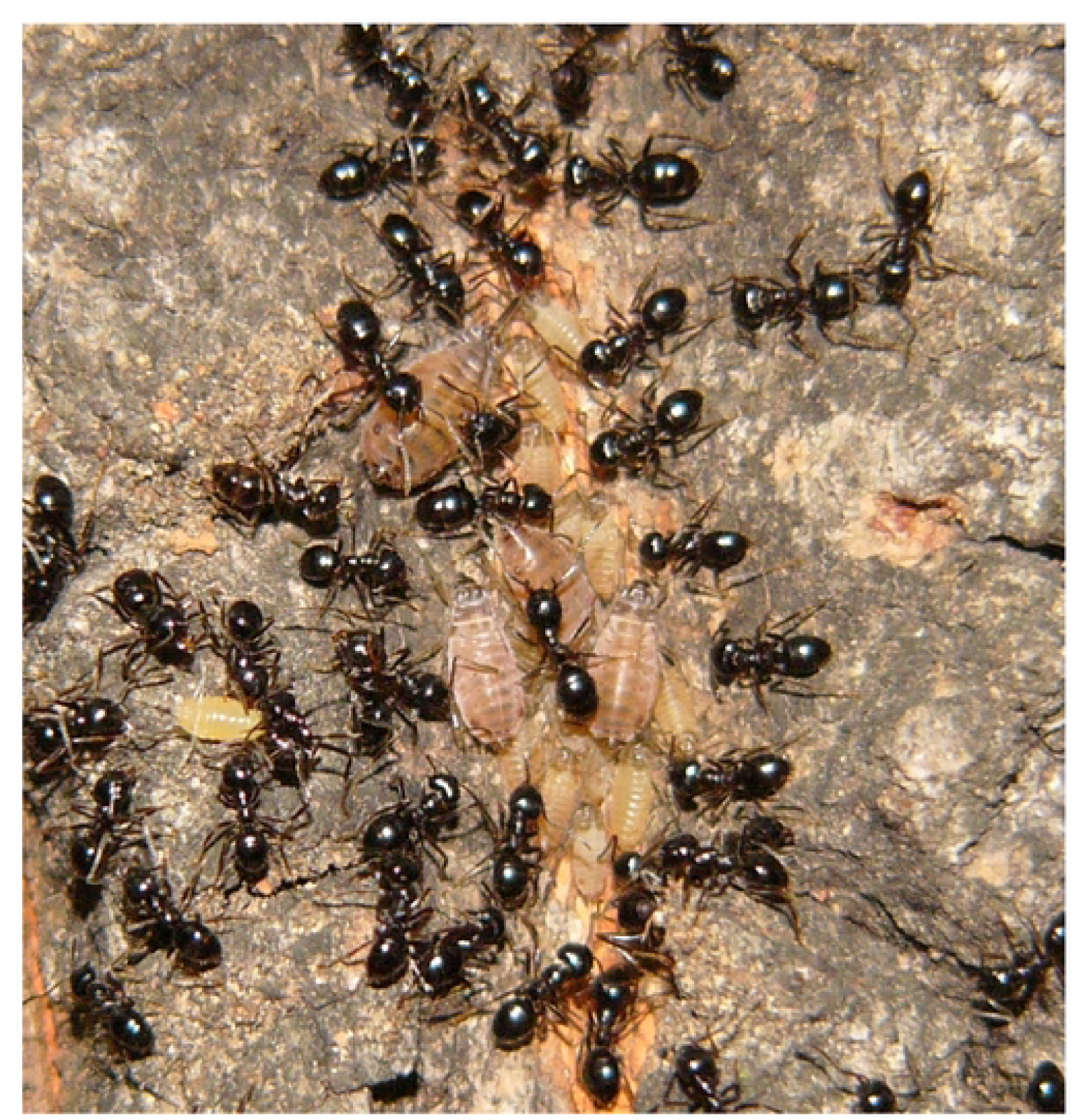
*Stomaphis japonica* colonies associated with *Lasius capitatus* ants on a *Quercus acutissima* tree trunk

## Material and Method

### Aphids and ants

*Stomaphis japonica* is one of the world’s largest aphids, growing to 7 mm in length. They form aggregations on the lower trunk surface of the Japanese chestnut oak *Quercus acutissima*, sucking phloem sap with proboscis (mouthparts) twice as long as their body length, and excreting honeydew [9]. *L. capitatus* is a jet black *Dendrolasius* (subgenus) ant with workers measuring around 4 - 4.5 mm long. Its biology is similar to that of its close relative *L. fuliginosus*, whose nests are usually found underground near tree roots, and whose workers travel along a conspicuous trunk trail between the nest site and aphids’ colony, both of which last for many years [10]. Ant attendance is required for completion of the aphids’ life-cycle because *Dendrolasius* ants transport young nymphs of *S. japonica* from the egg overwintering site at the base of trees to suitable feeding sites in the upper canopy of oak trees in spring [11].

### Field study

The study area was located in Matsumoto basin, Nagano, Japan. It is a mixed secondary forest of deciduous broad-leaved trees such as *Quercus acutissima, Quercus serrata*, and *Pinus densiflora*. We observed the presence of *S. japonica* aphids on 14 of the 22 *Q. acutissima* trees in 2004 (at the garden of Shorinji temple in Matsumoto), and on 40 of the 345 *Q. acutissima* trees in 2022 and 2023 (in a broader research area in Matsumoto). On a single host tree trunk, about 30–500 phloem-sap feeding aphid individuals are observed in summer in 2022 and 2023. Observations of ant predation was made on the one (in 2004) and two (in 2022-23) *Q. acutissima* trees where predation occurred frequently.

### Observation of attacks and predation by ants on aphids

We defined “attack” as the behaviors such as biting and application of formic acid by ants to aphids, and “predation” as the behaviors such as killing and transporting aphids to the ant nest. At Shorinji Temple in the suburbs of Matsumoto, Nagano, Japan (36.244651 N, 137.961440 E), we recorded the number of *S. japonica* aphid individuals up to a height of 1.6 m on a *Q. acutissima* tree trunk and measured the frequency with which predation occurred every 20 days from May 24 to October 6, in 2004. Each survey was conducted over a period of one to three days. *Lasius* ants were active throughout the day and night [12–14]. *Stomaphis* aphids have very low mobility and did not move from the same position on the tree trunk surface for as long as 60 hours [15]. Based on these observations, the number of predations per hour was multiplied by 24 to calculate the number of predations per day. Through our observations, we also observed parasitism by the predatory parasitoid wasp *Protaphidius nawaii* as another cause of mortality besides predation by ants. We also counted the number of mummies after aphid parasitism. Additionally, the body size of predated aphids was measured.

We observed several *S. japonica* colonies within a 7 km radius area near Futaba, Matsumoto, Nagano, Japan (36.210393 N, 137.962641 E), during August 1–31, 2022 and 2023. We observed a total of 39 *Q. acutissima* trees inhabited by *S. japonica*. There were five trees where predation of at least one individual aphid was observed over 3 hours of observation for each colony. We recorded the number of *S. japonica* aphid individuals on two of these five trees with a high frequency of predation by ants to determine how often predation occurred for a minimum of 10 hours.

### Manipulation experiment on honeydew excretion and ant predation

We investigated how honeydew excretion by aphids is effective in preventing ant attacks. Normally, ant predation begins with an attack on aphids, followed by an accelerated attacks on aphids by the several surrounding ant workers. To cause the ants to be aggressive and attack a particular aphid, we applied certain amount of ant secretion (including formic acid) to the aphid. First, we approached the fusiform brush to the ants. All of the ants attacked the brush without exception by biting it or spraying it with formic acid. We applied the brush containing certain amount of ant secretions (including formic acid) to the abdomen of an adult aphid that was pulling its proboscis out on a tree trunk for the purpose of inducing an ant attack on a specific aphid. This application was conducted 10 seconds after the brash-contact with the ants. Aphids to which the secretion was applied were usually attacked by ants within 3 minutes. We checked whether the attacking ants licked the honeydew excreted by the aphid, and whether they continued the attack if they did or did not lick the honeydew. We observed the attacking ants until they either stopped attacking the aphid and left or brought the aphid back to the nest. Although the ant secretions applied to the aphid may usually elicit a defensive behavior of ants, they served, in this experiment, as the substances that induce ants’ to the aphid. Ants’ aggression usually ceased when they licked the honeydew excreted by the aphid as shown in the results.

### Establishment of clonal lineages

We collected aphids at several distant localities in Nagano, Japan, from June 1 to June 30 in 2006, one adult female and her parthenogenetically produced larvae aphids on a *Q. acutissima* tree at each locality. We considered the aphids collected at each different locality to be separate clone because they were collected at localities at least 5 km away from each other. To establish clonal lineages, we attached 15 plastic cages to each of the two *Q. acutissima* tree trunks, introduced one clone individual into each of the cages, and reared it to adulthood allowing it to produce larvae. No ants were allowed to go into the cages and to attend the aphids. This procedure was also applied to the other clones of aphids collected from various locations, and finally seven clones were established. Each of the 7 clones were then randomly transferred to 30 cages set on the trunks of two *Q. acutissima* trees. We allowed them to reproduce, and thinned out the number of single clones in each cage during the experiment so that there were no more than 20 individuals.

### Estimation of the broad-sense heritability in honeydew excretion frequency

We counted the number of times honeydew was excreted by each individual of the seven clones. Each aphid was identified by applying acrylic paint to the dorsal surface of the abdomen. The presence or absence of paint did not affect the behaviour of aphids. We recorded the number of times they excreted honeydew once a day in 10 minutes for an individual, ranging from 1 to 10 individuals in a single cage. Observations were made between 8:00 AM and 2:00 PM.

Only aphid individuals for which data were obtained for at least 3 days (10 minutes x 3 times) per instar stage (3rd, 4th and adult stages) were analysed for the frequency of honeydew excretion. Clones were analysed only when data were obtained for nine or more individuals of the clone. We averaged the number of times of ten-minutes’ nectar excretion for each individual (for each instar stage) on the more-than-three days of ten-minutes’ measurement, and used this as the data for the frequency of honeydew excretion for that individual (for each instar stage). Since the ratio of the standard deviation to the mean of the frequency of honeydew excretion, i.e., the coefficient of variation, exceeded 0.2, we added 1 to the original data (the frequency of ten-minutes’ nectar excretion) and used the Log10 transformed value for the calculation of genetic variance and heritability.

We estimated genetic variance (*V*_*G*_) and environmental variance (*V*_*E*_) by a one-way Type II (variate model) analysis of variance. In our study, the mean square of the measurements within a clone (*MS*_*within*_) at each instar stage is the estimate of the environmental variance (*V*_*E*_) [16]. On the other hand, the mean squares among clones (*MS*_*groups*_) are the environmental variance plus the variation among clones as follows [17].

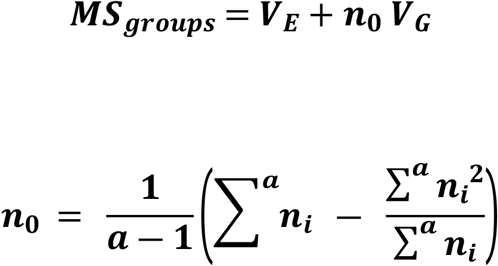

Where *n*_*i*_ is the number of observations for *i*th clone and *a* is the number of clones. Based on the above, *V*_*G*_ is as follows.

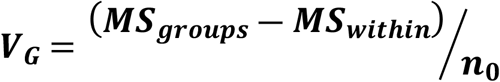

We estimated heritability in the broad sense as the ratio of genetic variance to the sum of genetic and environmental variance. We tested for differences in the frequency of honeydew excretion between clonal lines in the same cage by ANOVA.

## Result

*S. japonica* aphids were heavily preyed upon by *L. capitatus* (Fig. 3). On 20 July and 13 September, ants are estimated to have eaten about half of the aphids on the *Q. acutissima* tree per day (Fig. 3). Similarly, during August 1–31, 2022 and 2023, predation by ants occurred at a rate of about 5.4 individuals per hour (279 individuals / 51.5 h, that is c.130 individuals / day) on two *Q. acutissima* trees where frequent ant-predation occurred. As each of the two trees had 200–300 aphids in August, ants are estimated to have eaten more than half of the aphids on the two *Q. acutissima* trees per day. In comparison, parasitism by *Protaphidius nawaii* wasps, which is the only natural enemies of *S. japonica* detected in the study area, was relatively low, at a rate of about 0.08 individuals per hour (231 individuals / 6 day).

**Fig. 3.**
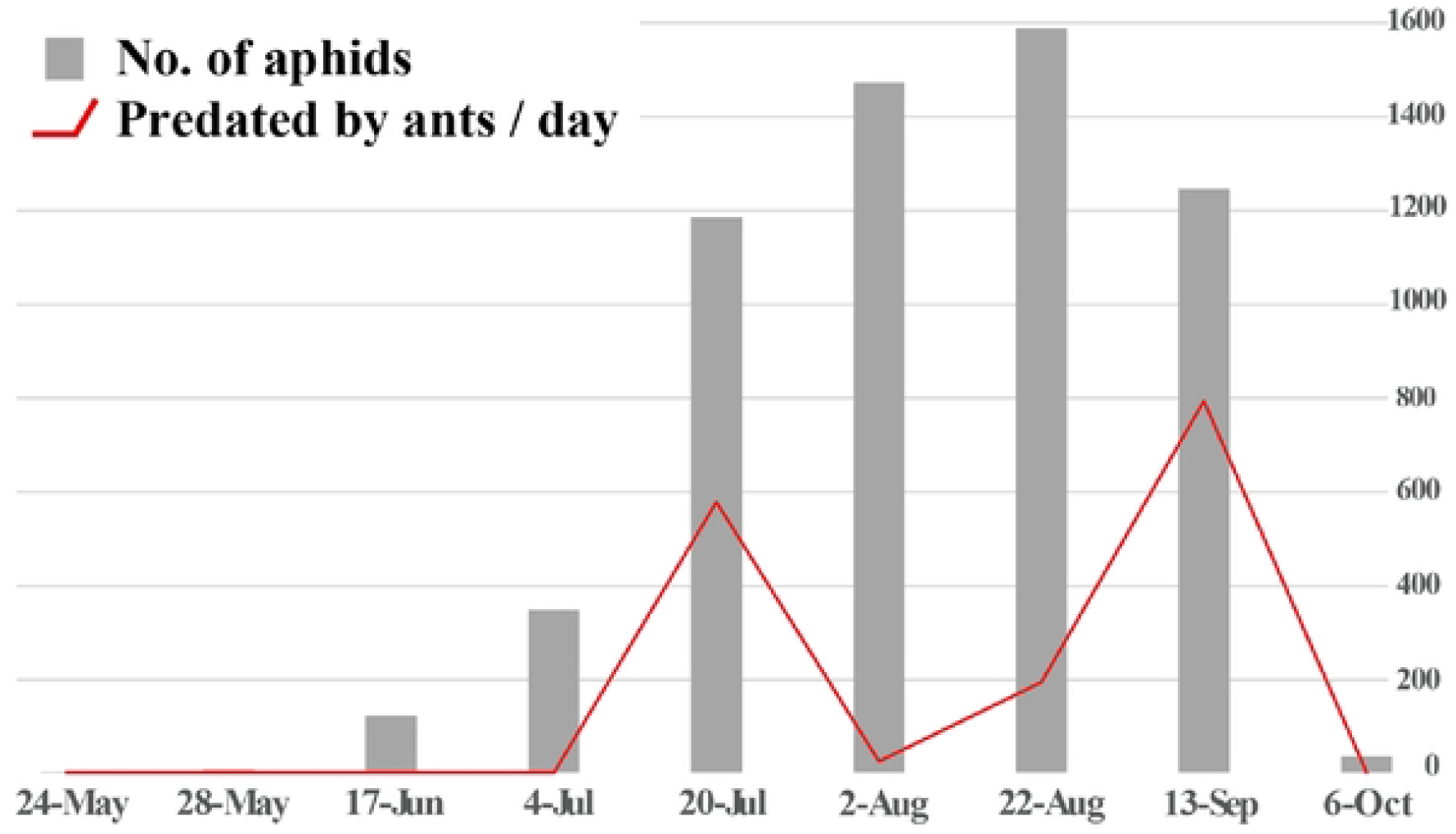
Seasonal fluctuations in *Stomaphis japonica* aphid population size and the number of aphids predated by *Lasius capitatus* ants on a *Quercus acutissima* tree

The ants selectively preyed on the aphids which did not provide honeydew (Table 1, Fig. 1). An attacking *L. capitatus* tends to stop attacking when it received honeydew from the target aphid, while it tends to continue attacking and finally eat the aphid unless it did not receive honeydew from the target aphid (Table 1; Fisher’s exact test *P*<0.001).

**Table 1.**
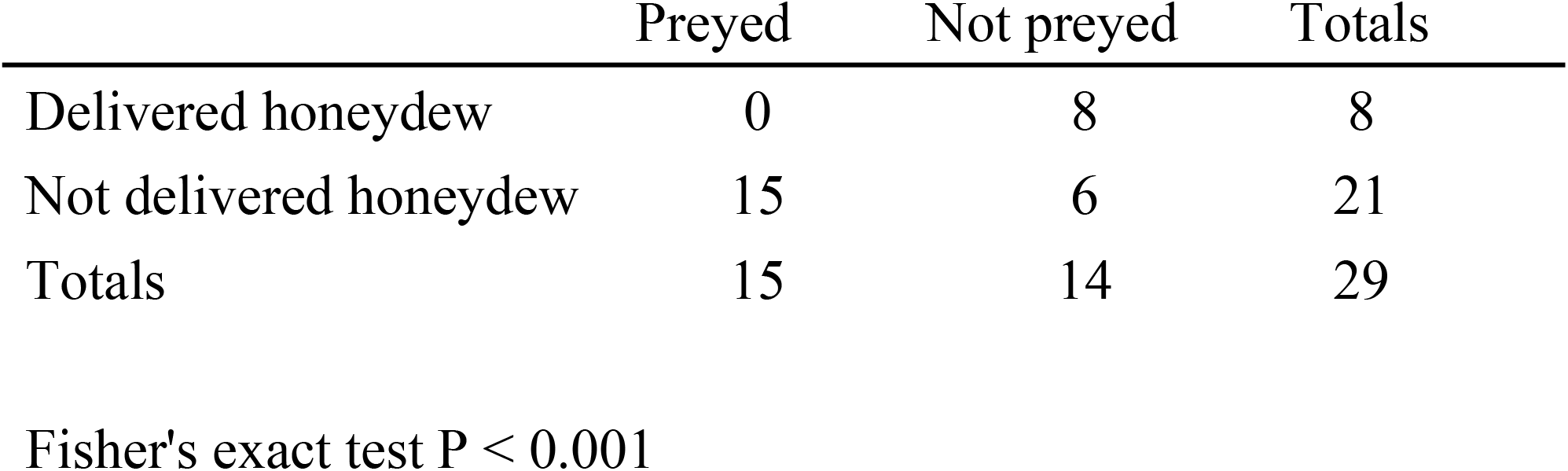
Effect of honeydew delivery on aphid survival from ant predation.

|  | Preyed | Not preyed | Totals |
| --- | --- | --- | --- |
| Delivered honeydew | 0 | 8 | 8 |
| Not delivered honeydew | 15 | 6 | 21 |
| Totals | 15 | 14 | 29 |

Fisher's exact test $P < 0.001$

The heritability in the broad sense for the frequency of honeydew excretion per 10 minutes was 0.098, 0.085 and 0.068 for the third instar larvae, forth instar larvae and adults, respectively (Table 2). The frequency of honeydew excretion was significantly different among clonal lines in 3rd instar larvae (ANOVA, Sum of Square = 1.185, Mean Square = 0.296, *F*-value = 2.890, *P* = 0.027), close to significant in 4th instar larvae (Sum of Square = 0.863, Mean Square = 0.216, *F*-value = 2.296, *P* = 0.066) and not significant in adults (Sum of Square = 0.354, Mean Square = 0.177, *F*-value = 2.080, *P* = 0.137).

**Table 2.** Genetic variance, environmental variance, and broad-sense heritability calculated for the number of times of honeydew excretion per 10 minutes.

|  | 3rd instar | 4th instar | Adult |
| --- | --- | --- | --- |
| genetic variance | 0.011 | 0.009 | 0.006 |
| environmental variance | 0.103 | 0.094 | 0.085 |
| heritability | 0.098 | 0.085 | 0.068 |

## Discussion

Our result suggests that aphids should deliver honeydew to survive when being attacked by ants (Table 1, Fig. 1), and frequency of honeydew excretion has genetic basis (Table 2). Consequently, the aphids which cannot excrete honeydew when attacked by ants are prone to be predated. As the predation pressure by ants is sometimes very high (Fig. 3), the aphids with low honeydew secretion are susceptible to predation by ants.

Honeydew quality and quantity is important in the maintenance of and competition for mutualistic services of ants [5, 18, 19]. Honeydew production is also reported to be important for escaping ant predation. When the aphid *Lachnus tropicalis* is attacked and excretes honeydew which is consumed by the attendant ant *Lasius niger*, the predation rate is significantly lower than in cases where the aphids fail to yield honeydew to the ants [4]. In addition, *L. niger* has been shown to prey preferentially on aphid species which produce less honeydew [20]. Here, we found a slight genetic basis for the frequency of honeydew excretion of *S. japonica* (Table 2). This suggests frequency of honeydew excretion can evolve; the more frequency of honeydew excretion, the higher the probability of nectar being passed on to ants, and the lower the probability of the aphid being preyed on. In fact, since there are 5–7 clonal lineages in the *S. japonica* population on a single *Q. acutissima* tree under natural conditions (T. Matsuura, unpublished observation), it is possible that ants encounter variation in the amount of honeydew secretion among clones under natural conditions.

Aphids that have longer proboscis should have an advantage in procuring phloem sap deeper in the tree stem, and thus excrete honeydew easily. Thus, we can assume that selective predation by ants will extend proboscis length of their aphid livestock. Indeed, within the aphid genus *Chaitophorus*, ant-tended species have longer probosciss than untended species [21]. The selection, however, appears not so extreme in most aphid species because they can escape from the ant attacks in the long run by fling. The point is that most aphid species are diffusely associated with ants (one species of aphid is associated with many genera of ants), and they are not in captivity, with their population size not being regulated by particular ant species. In contrast, the *Stomaphis-Dendrolasius* association has a highly specific, dependent and captive nature. The aphid population is well regulated by massive predation of the ants (Fig. 3), and their colony persist over the years tended by a single ant colony with the overwintering eggs being laid on the tree trunk (TM, personal observation).

After all, this study found that predation by *Dendrolasius* ants is selective, *Stomaphis* aphids with low amounts of honeydew tend to be more susceptible to predation, and the frequency of honeydew excretion by aphids has a genetic basis. We anticipate that other cases of selective predation on mutualists with preferred characteristics will be found in nature, especially in obligate and captive associations in terrestrial and aquatic systems, and that in-depth research on the cases will bring new horizons to ecological and evolutionary science.

## Ethics

This study did not require ethical approval from a human subject or animal welfare committee.

## Data accessibility

All data concerning this study are available upon appropriate request by e-mailing T.I. or T.M.

## Authorship contributions

Matsuura T. drafted the manuscript. Handa C., Matsuura T. and Takahashi S. carried out the field investigation, conducted the statistical analysis and critically revised the manuscript. Itino T. designed the study and drafted the manuscript. All authors gave final approval for publication and agree to be held accountable for the work performed therein.

## Acknowledgments

We thank S.-P. Quek, D. Hembry and U Ban for editorial assistance and comments on the manuscript. We thank the residents living in the study area for their understanding of this study. We thank S. Duhon for English editing.

